# Genome-scale prediction of gene ontology from mass fingerprints reveals new metabolic gene functions

**DOI:** 10.1101/2024.11.27.625594

**Authors:** Christopher J. Vavricka, Masao Mochizuki, Satoshi Yuzawa, Masahiro Murata, Takanobu Yoshida, Naoki Watanabe, Masahiko Nakatsui, Jun Ishii, Kiyotaka Hara, Hal S. Alper, Tomohisa Hasunuma, Akihiko Kondo, Michihiro Araki

## Abstract

Mass-based fingerprinting can characterize unknown strains, however expansion of these methods to predict specific gene functions is lacking. Therefore, rapid mass fingerprinting was developed to functionally profile a comprehensive yeast knockout library. Matrix assisted laser desorption ionization (MALDI)-time of flight (TOF) mass fingerprints of 3,238 Saccharomyces cerevisiae knockouts were digitized for correlation with gene ontology (GO) annotations. Random forests and support vector machine (SVM) algorithms precisely assigned GO accessions with AUC scores all above 0.83. SVM was the best predictor with average true positive and true negative rates of 0.975 and 0.991, respectively. The SVM model suggested new functions for 28 uncharacterized yeast genes. Metabolomics analysis of two knockouts (YDR215C and YLR122C) of uncharacterized genes predicted to be involved in methylation-related metabolism, showed altered intracellular contents of methionine-related metabolites. Increased S-adenosylmethionine in YDR215C highlights potential for enhancement of methylation pathways. These results demonstrate that MALDI-TOF fingerprints can be rapidly digitized, resulting in datasets that enable prediction of microbial genotypes and even the function of specific genes. This fingerprinting method can inform optimal bioproduction chassis selection.

## Introduction

Many recent advances in ‘omics’ methods have attempted to map global views of various cells. However, it is still expensive and time-consuming to apply these approaches to analyze libraries of engineered microorganisms. The combination of artificial intelligence with synthetic biology offers potential to speed up the functional prediction of individual strains in microbial libraries (1-4).

Mass spectra are especially convenient to process for machine learning analysis and have even been applied to the discrimination of microbial and human cell populations (5-8). One of the first examples of an artificial intelligence application to scientific research is the Dendral system, which was designed to determine chemical structures from mass spectra (9). In addition, rapid mass spectrometric analysis, as well as Fourier transform infrared spectroscopy analysis, can be used to generate fingerprints for microbial strains, and machine learning models have been reported to discriminate species based on the fingerprints (10-15). However, no previous fingerprint studies that we are aware of have comprehensively characterized an entire gene knockout library to gain insight into specific gene functions.

While identifying the function of unknown genes has been greatly accelerated by new machine learning models including DeepEC (16), CLEAN (17) and EnzymeNet (18, 19), these models only predict the functions of enzyme; however, many currently unknown genes are not enzymes. Furthermore, the mentioned supervised learning models require labeling of the data based on enzyme database annotations which are often incorrect or lacking for unknown genes.

Therefore, the current study aims to demonstrate that matrix assisted laser desorption ionization (MALDI)-time of flight (TOF)-based mass fingerprints can be digitized to rapidly establish a large dataset, and then the dataset can be mined to predict the function of microbial strains and even individual genes. This rapid fingerprinting strategy can enable optimal bioproduction chassis selection, without requiring tedious targeted analyses of the entire genome, metabolome or proteome (20, 21).

Compared to other fingerprinting analysis methods, MALDI-TOF is more rapid and convenient. MALDI-TOF fingerprinting does not require a cell lysis or extraction step; the cells can be directly taken from the culture and dropped directly onto the MALDI analysis plate. With the ability for increased-throughput, a high-throughput MALDI-TOF-based diagnostic tool can more rapidly estimate functional changes of cells under different conditions, or predict genotypes of variants from strain libraries, powerful tools for the optimization of cell performance.

In this study, the yeast knockout library of the *Saccharomyces* Genome Deletion Project was selected to develop a high-throughput method to predict genotype and gene function from MALDI-TOF mass fingerprints. As a model eukaryotic organism, *Saccharomyces cerevisiae* has great potential as a cell factory chassis, and many advantages in terms of fermentation, genetic manipulation, protein processing and availability of comprehensive omics data (20-23). According to the Saccharomyces Genome Database (https://www.yeastgenome.org/genomesnapshot) 10% of *S. cerevisiae* genes are uncharacterized and an additional 10% of genes are classified as ‘dubious’. Therefore, *S. cerevisiae* was selected to develop methods for rapid genotype prediction from mass spectrometric data. Previously, machine learning has been applied to the analysis of various wine and brewing yeast strains (24), however there are no reports on the analysis of a comprehensive knockout library using MALDI-TOF fingerprints and machine learning.

High quality MALDI-TOF fingerprints were obtained from total cell extracts of 3,238 *S. cerevisiae* single gene knockout strains. Several machine learning models were developed to correlate the mass fingerprints with yeast gene ontology (GO) annotations. Support vector machine (SVM) models could quickly and precisely assign GO accessions to yeast knockouts with AUC scores above 0.83. Average AUC scores for SVM and random forests prediction were 0.991 and 0.981, respectively. This new approach offers high potential for the rapid characterization of strains with unknown genotype. Accordingly, the SVM models were able to suggest functions for 28 uncharacterized genes, which have remained uncharacterized since at least 2019. Further metabolomics data were consistent with the predictions for two selected knockout strains.

## Results and Discussion

### High-throughput MALDI-TOF analysis of yeast deletion mutants

The comprehensive library of 4,847 *S. cerevisiae* knockouts was obtained from Invitrogen and maintained on 96 well plates. Automatic high-throughput yeast cell extraction with formic acid was performed on the plates as described in the methods section.

α-Cyano-4-hydroxycinnamic acid (HCCA), sinapinic acid (SA) and 2,5-dihydroxybenzoic acid (DHB) were first tested as matrix with wildtype yeast extract (Figure 1). Although SA and DHB had a lower frequency of quality raster spots, SA performed best in terms of compatibility with automatic measurement, uniform distribution of spot crystals, narrow peak width and better quality high molecular weight peaks. Therefore, SA was selected for high-throughput MALDI-TOF analysis.

**Figure 1.**
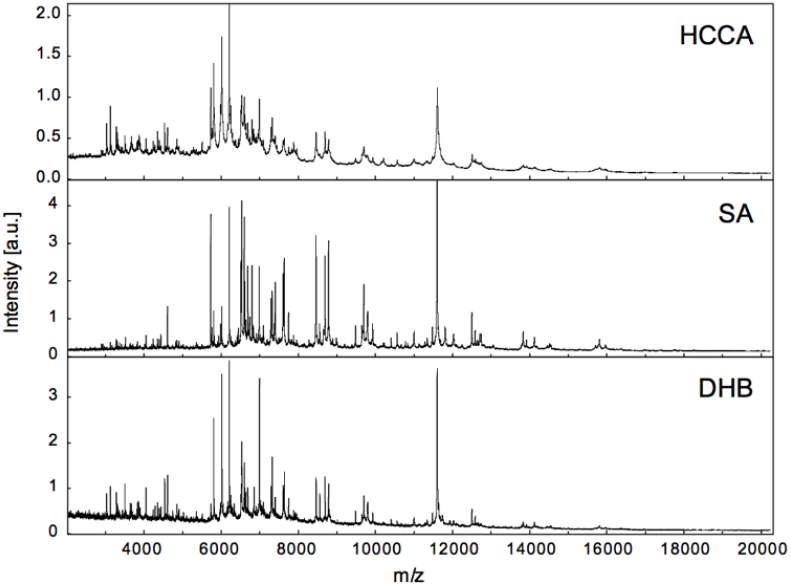
Comparison of *S. cerevisiae* total MALDI-TOF spectra using α-Cyano-4-hydroxycinnamic acid (HCCA), sinapinic acid (SA) and dihydroxybenzoic acid (DHB) as matrix.

Loss of an ion peak may indicate the loss of a corresponding protein encoded by a knocked-out gene (Figure 2). To enable automatic comparison of peaks between mass spectra, all spectra were converted to binary vectors. For each MALDI-TOF spectra, a mass window of *m/z* 3,000 to 20,000 was divided into 1,700 segments at intervals of 10 *m/z* units for processing into 1,700-digit binary vectors. The window of *m/z* 3,000 to 20,000 was selected due to the presence of high noise and baseline below *m/z* 3,000, and the upper mass limit of the MALDI-TOF instrument, respectively. The resulting vectors were then used for gene ontology (GO) prediction (Figure 3).

**Figure 2.**
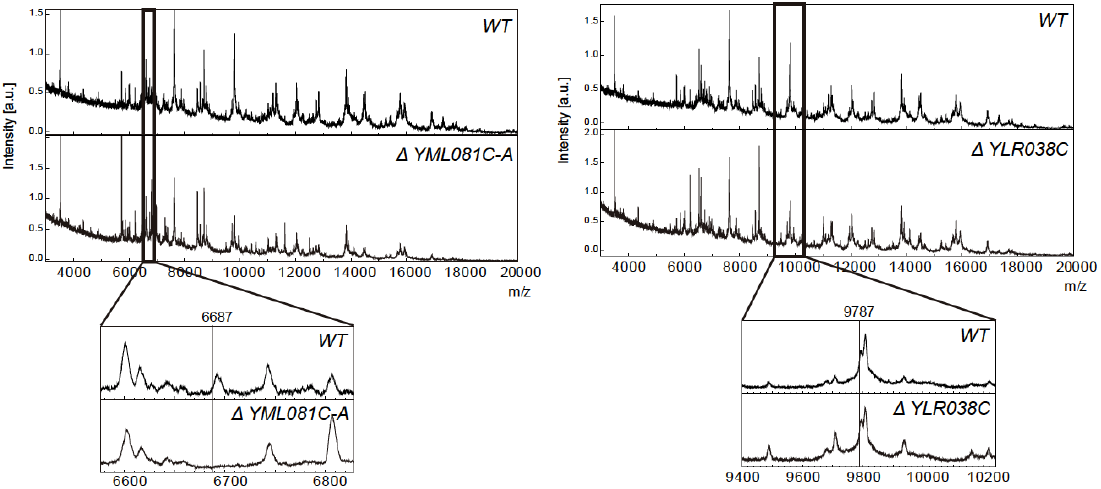
Knockout of MALDI-TOF peaks corresponding to yeast gene products. In a few spectra, the knockout of a gene product could be observed, as was the case for the ATP18 subunit of the mitochondrial F_1_F_0_ ATP synthase (left panel). However, in most spectra, no change could be observed in the predicted *m/z* region corresponding to the respective gene product, as shown for knockout of COX12 (right panel).

**Figure 3.**
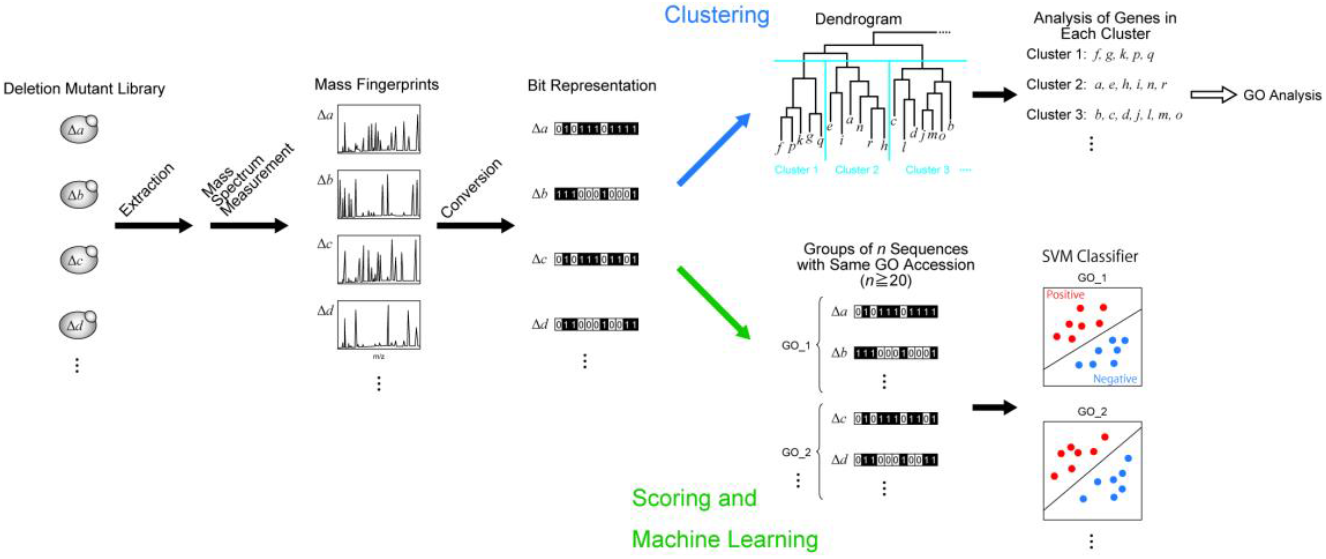
Processing of MALDI-TOF spectra for the prediction of gene ontology (GO).

### Computational prediction of GO accessions

After a preliminary clustering analysis to check if the digitized mass fingerprints correlated with GO accessions (Supplementary Figure 1), we hypothesized that fingerprints from the same GO family will share some similar features that are difficult to see by eye; therefore, computer algorithms might be able to infer potential elusive features from the dataset. This approach offers potential to infer the functions of variant genes from various microbial strain libraries.

Accordingly, Tanimoto (25, 26), random forests (27) and SVM (28) models were built to accurately predict the GO relationships. A total of 3,238 yeast gene knockout strains were included in the mass fingerprinting and computational analyses, according to the methods section. This dataset represents 66.8% coverage of the entire 4,847 gene knockout library. Some loss of coverage was due to quality control of inadequate spectra resulting from a heavy workload of the detector.

### Cross validation of GO predictions

Cross validations were performed to provide unbiased evaluations of each prediction method (Figure 4). Of the three methods, the Tanimoto scoring method was the least accurate GO predictor (Figures 4 and 5). When performing the analysis on the 1,559 GO accessions matching 3 or more binary vectors, the Tanimoto model performed moderately. However, when using only the 332 GO accessions matching 20 or more binary vectors, the Tanimoto algorithm was unreliable, with 126 GO accessions producing indistinguishable positive and negative Tanimoto distributions, 206 GO accessions producing partially overlapping Tanimoto distributions and no GO accessions producing Tanimoto distributions with 95% separation (Figure 5). This indicates that simple scoring methods are not sophisticated enough for reliable GO prediction, and therefore machine learning algorithms were next tested.

**Figure 4.**
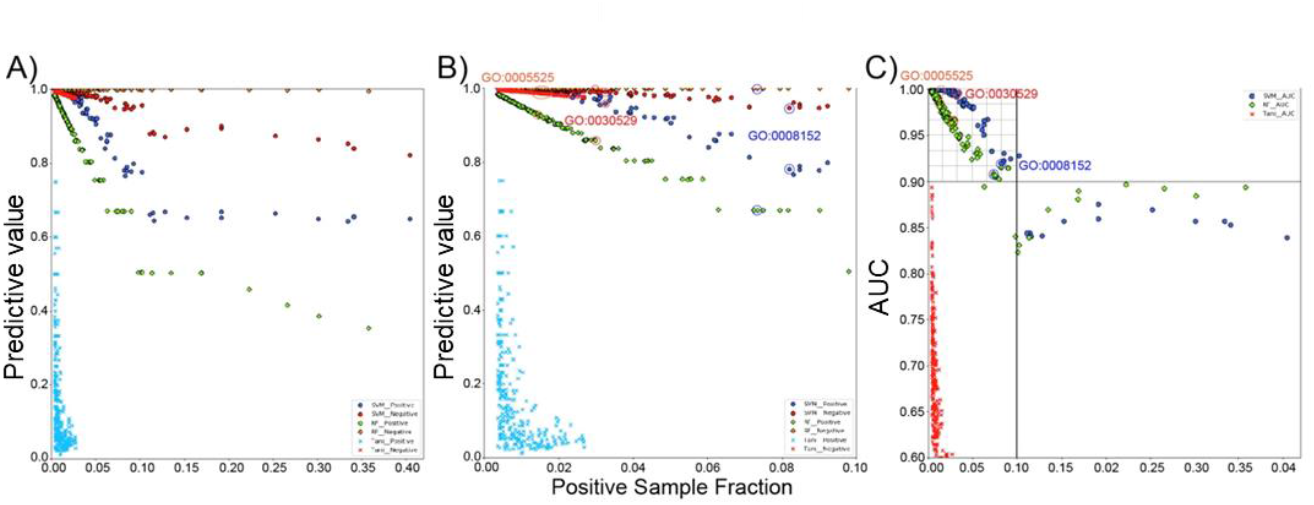
Cross validation of GO accession prediction models. **(A)** Positive and negative predictive values for each prediction model. (**B**) Predictive values for GO accessions with positive sample fractions up to 0.10. Positive predictive values were calculated by dividing the number of true positives by the sum of the true positives and false positives. Negative predictive values were calculated by dividing the number of true negatives by the sum of true negatives and false negatives. Blue and red circles represent positive and negative predictive values of SVM prediction, respectively. Green and orange diamonds represent positive and negative predictive values of random forests prediction, respectively. Blue and red cross marks represent positive and negative predictive values of Tanimoto prediction, respectively. **(C)** Area under the curve (AUC) scoring for Tanimoto, random forests and SVM models. Increase in the positive example sample size by SMOTE (synthetic minority over-sampling technique) may explain why scores stop decreasing around 0.1 sample ratio. Random forest and SVM results for GO:0005525 (GTP binding, orange), GO:0030529 (ribonucleoprotein complex, red) and GO:0008152 (metabolic process, blue) are presented in panels **B** and **C**. Blue circles, green diamonds and red cross marks represent AUC scores of SVM, random forests and Tanimoto prediction, respectively.

**Figure 5.**
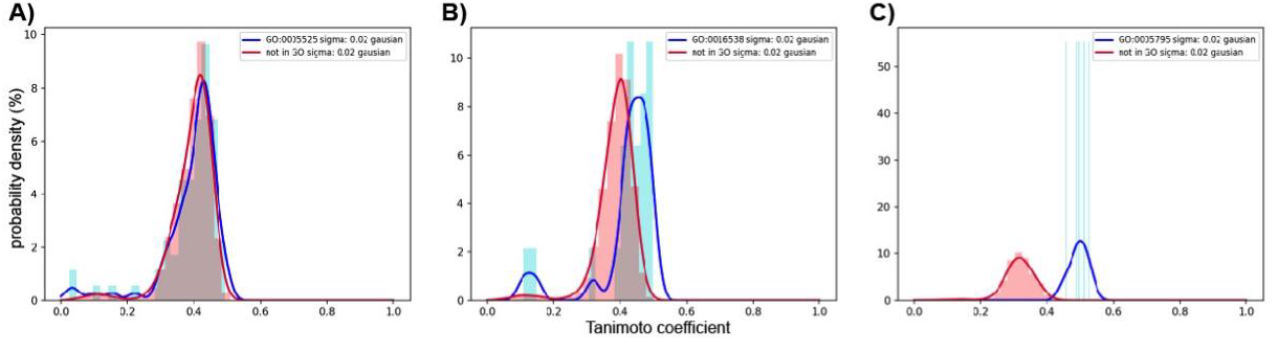
Tanimoto probability density distributions for 3 of the 1,559 GO accessions matching 3 or more binary vectors. **A)** Tanimoto distribution for GO accession GO:0005525 (GTP binding) with positive matching to 75 of the 5,356 binary vectors (0.014 positive matching ratio). 266 GO accessions produced indistinguishable Tanimoto curves that could not be separated with 95% confidence with a Mann–Whitney *U* test. **B)** Tanimoto distribution for GO accession GO:0016538 (cyclin-dependent protein serine/threonine kinase regulator activity) with positive matching to 24 binary vectors (0.0045 matching ratio). 958 GO accessions produced partially overlapping Tanimoto curves that could be separated with 95% confidence with a Mann–Whitney *U* test. **C)** Tanimoto distribution for GO accession GO:0005795 (Golgi stack) with positive matching to 5 binary vectors (0.00093 matching ratio). 335 GO accessions produced Tanimoto curves with 95% separation. Positive distribution curves are drawn in blue and negative distribution curves are red.

As our data set was sufficiently large and can be continuously expanded, machine learning methods are much more attractive than simple scoring methods such as the Tanimoto correlation. Accordingly, random forests was able to predict GO accessions with an average AUC score of 0.981, and an average true positive rate of 0.925. Interestingly, the random forests method resulted in near perfect true negative rates of 0.999 (Figures 4 and 6). In Figure 6, overlap of matching and non-matching distributions is observed for random forests positive prediction representing false positives, while almost no false negatives are observed. Cross validation of the random forests predictions resulted in a false negative rate of 0.003 and a false positive rate of 0.33 when the number of binary vectors in each GO accession group was between 20 and 500. However, for high sample GOs, the false positive rates were often 0.5 or more, a level too high to use practically.

**Figure 6.**
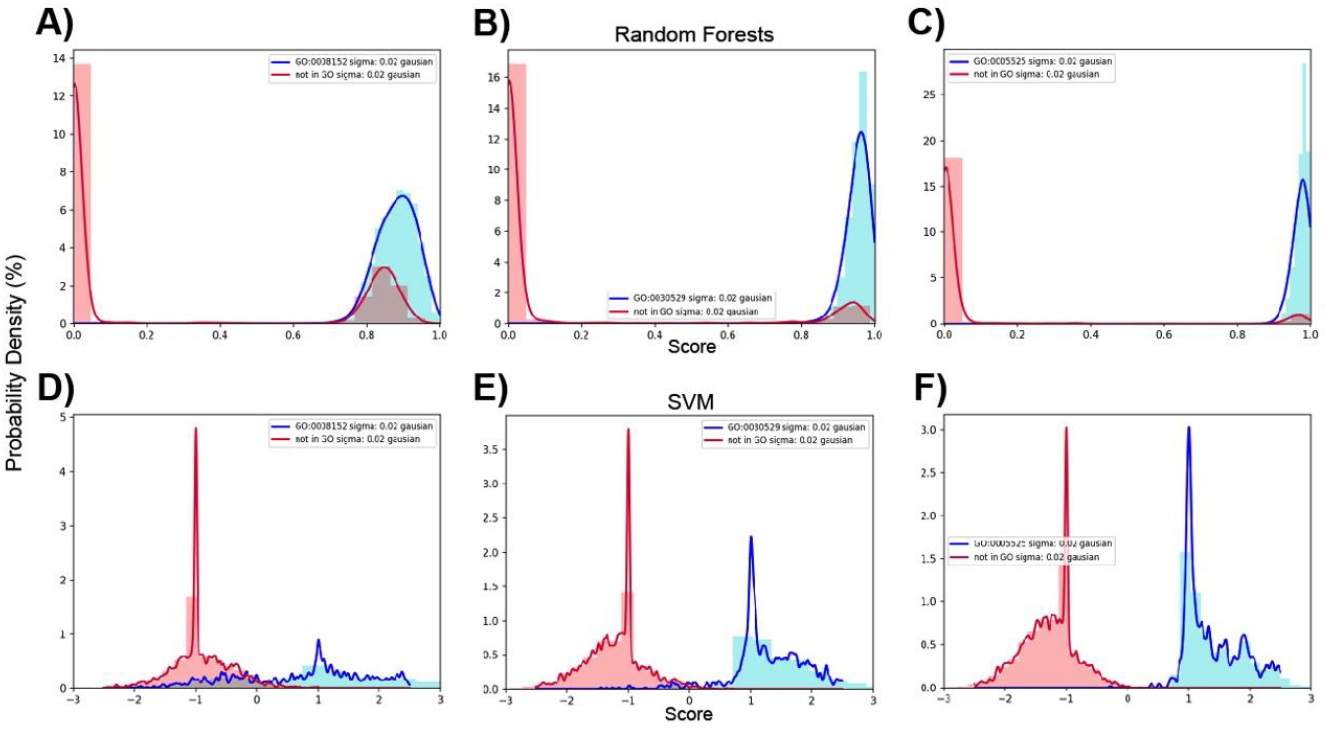
Random forests and SVM score density distributions for 3 of the 332 GO accessions matching 20 or more binary vectors. **A)** Random forests distribution for GO accession GO:0008152 (metabolic process) with positive matching to 393 of 5,356 binary vectors (0.073 positive matching ratio). Random forest scores are given as the probability of positive samples. **B)** Random forests distribution for GO accession GO:0030529 (ribonucleoprotein complex) with positive matching to 159 binary vectors (0.0297 positive matching ratio). **C)** Random forests distribution for GO accession GO:0005525 (GTP binding) with positive matching to 75 binary vectors (0.014 positive matching ratio). **D)** SVM distribution for GO:0008152 (metabolic process) with positive matching to 393 of 4,731 binary vectors (0.083 positive matching ratio). SVM scores represent distances of binary vectors from the decision boundary and are given as a multiple of the margin distance. **E)** SVM distribution for GO:0030529 (ribonucleoprotein complex) with positive matching to 159 binary vectors (0.034 positive matching ratio). **F)** SVM distribution for GO:0005525 (GTP binding) with positive matching to 75 binary vectors (0.016 positive matching ratio). Positive prediction distributions are drawn in blue and negative distributions are red.

Compared to the above Tanimoto scoring and random forests models, SVM was the best predictor of GO accessions with an average AUC score of 0.991 and an average true positive rate of 0.975. In contrast to the random forests results, overlap of distributions is apparent for SVM negative prediction representing some false negatives, while few positives are observed (Figure 6). SVM negative prediction was lower than that of random forests, but the SVM average true negative rate was still very high at 0.991. For GO accessions associated with a lower pool of binary vectors (20 - 65 binary vectors), the SVM false positive rate was 0, which may be due to overfitting. Despite these minor tradeoffs, the SVM model was the best overall method for predicting GO accessions (Figures 4 & 6), offering a new empirical method to predict the function of uncharacterized genes.

### Real prediction of gene ontology for genes with unknown functions

In our prediction models, knockout strains of genes annotated as uncharacterized or dubious would be considered as false positives when matching to specific GO accession group; however, we realized that some of these matches may infer the actual function of these uncharacterized genes. Therefore, the random forests and SVM models were tested against the data of knockouts with unknown function (Figure 7).

**Figure 7.**
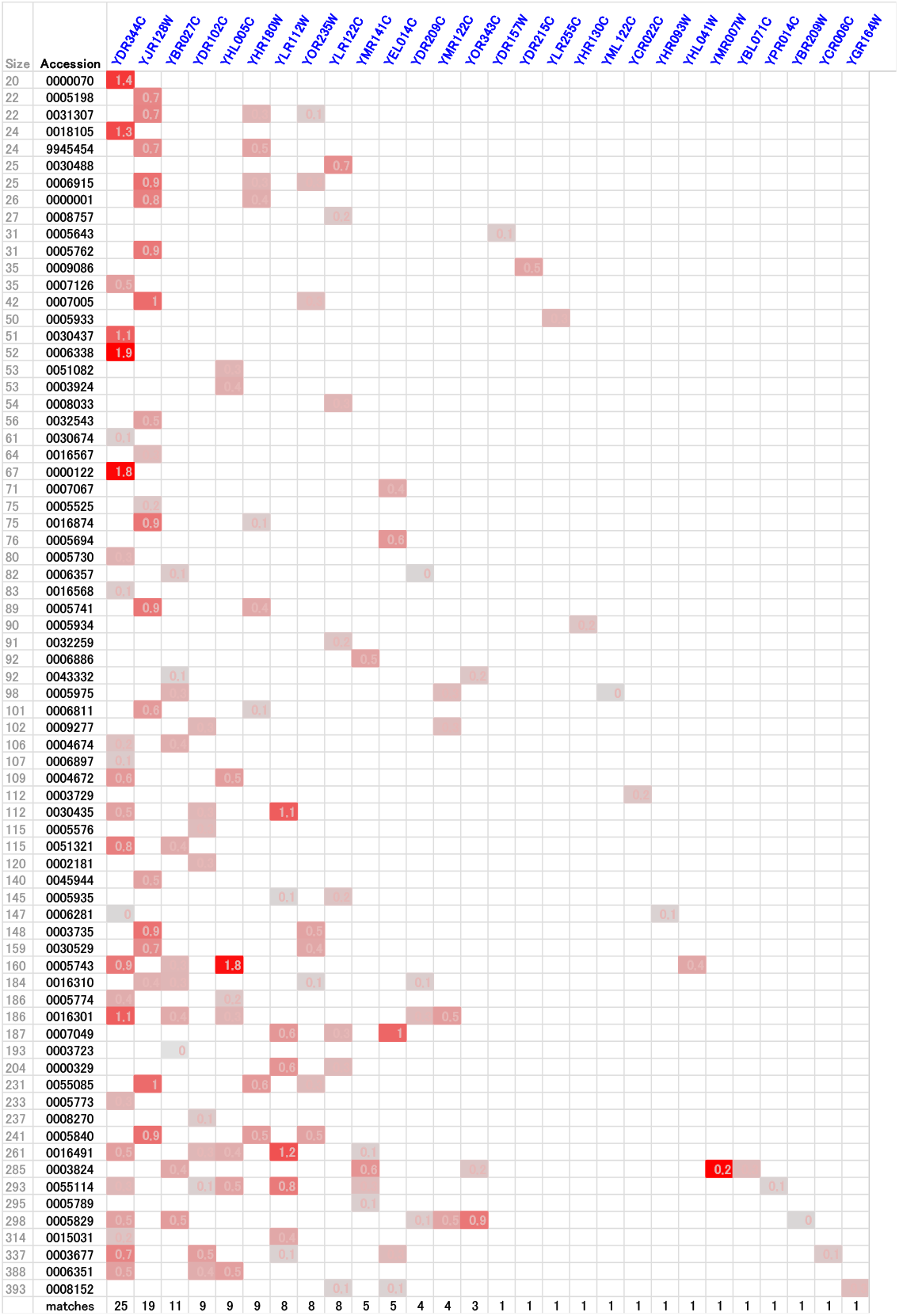
Heatmap of SVM-based GO matching to unknown yeast gene knockouts. YDR344C, YJR128W, YBR027C, YDR102C, YHL005C, YHR180W, YLR112W, YOR235W, YLR122C, YMR141C, YEL014C, YDR209C, YMR122C, YOR343C, YDR157W, YDR215C, YLR255C, YHR130C, YML122C, YCR022C, YHR093W, YHL041W, YMR007W, YBL071C, YPR014C, YBR209W, YCR006C and YGR164W are all annotated as knockouts of uncharacterized genes in the Saccharomyces Genome Database (www.yeastgenome.org). For large GO accession groups that contain approximately 500 or more gene knockout vectors, matching was too high to be meaningful. Therefore, SVM-based matching to GO accessions which include less than 400 gene knockout vectors was analyzed for this heatmap. YDR215C matched to methionine biosynthetic process (GO:0009086) only. YLR122C matched to *S*-adenosylmethionine-dependent methyltransferase activity (GO:0008757), metabolic process (GO:0008152), methylation (GO:0032259), tRNA processing (GO:0008033), tRNA methylation (GO:0030488), cell cycle (GO:0007049), fungal-type vacuole membrane (GO:0000329) and cellular bud neck (GO:0005935) (Supplementary Figure 2). YBR027C matched to GO:0006357 (regulation of transcription by RNA polymerase II), GO:0043332 (mating projection tip), GO:0005975 (carbohydrate metabolic process), GO:0004674 (protein serine/threonine kinase activity), GO:0051321 (meiotic cell cycle), GO:0005743 (mitochondrial inner membrane), GO:0016310 (phosphorylation), GO:0016301 (kinase activity), GO:0003723 (RNA binding), GO:0003824 (catalytic activity) and GO:0005829 (cytosol).

Many of the GO accession groups which matched to a particular unknown knockout, were related to each other hierarchically (Supplementary Figure 2), suggesting that the predicted functions are meaningful. From the heatmap (Figure 7), two target gene knockout strains, YDR215C and YLR122C, were predicted to be involved in methylation-related metabolic pathways. YDR215C matched to the GO accession methionine biosynthesis. YLR122C matched to methylation, tRNA processing, tRNA methylation and *S*-adenosylmethionine (SAM)-dependent methyltransferase activity (Supplementary Figure 2). Due to our previous findings on the importance of methylation-related metabolic pathways on the bottleneck steps benzylisoquinoline alkaloid methylation (29-31), we decided to analyze these strains further.

Based on the SVM models, YDR215C and YLR122C were identified as potential yeast strains with altered methylation-related metabolism (Figure 7). Therefore, metabolomics analysis of these strains was performed. When grown in minimal medium, YDR215C and YLR122C were found to contain altered intracellular levels of SAM and methionine, relative to the wild-type strain (Figure 8, A & B). Differences in the nucleotide levels were also observed between the knockout and wild-type strains. Remarkably, YDR215C contained 5-fold higher levels of SAM and 1.8-fold higher levels of methionine compared to the wild-type strain. On the other hand, YLR122C contained approximately half the level of intracellular SAM and similar levels of methionine compared to that of the wild-type. Accordingly, SAM-dependent methylation of benzylisoquinoline alkaloids was also tested in these strains. While the observed differences in alkaloid methylation, relative to that of the wild-type strain, were slight, relative increases in coclaurine and NMC were observed in 4 out of 4 conditions for YDR215C and decreases were observed in 4 out of 4 conditions for YLR122C (Figure 8C); these results are consistent with the observed intracellular SAM levels.

**Figure 8.**
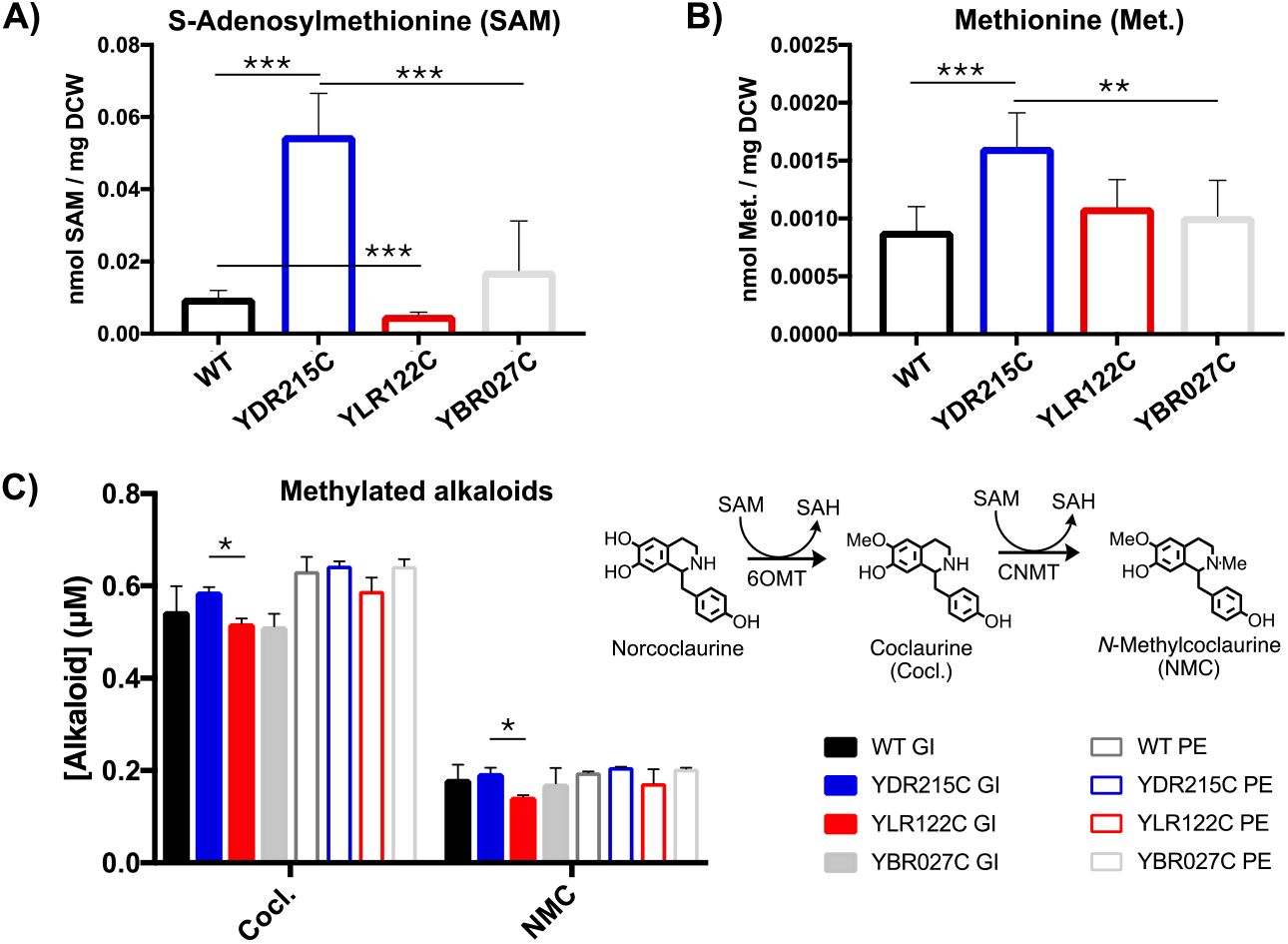
Yeast knockout YDR215C shows enhanced methylation metabolism. **A)** Intracellular SAM concentrations; for each condition two independent cultures were analyzed two times each for a total of four samples (n=4). **B)** Intracellular methionine (Met.) concentrations; for each condition two independent cultures were analyzed two times each for a total of four samples (n=4). **C)** Conversion of norcoclaurine to coclaurine (Cocl.) and *N*-methylcoclaurine (NMC) by norcoclaurine 6-*O*-methyltransferase (6OMT) and coclaurine *N*-methyltransferase (CNMT); for each condition two independent cultures were analyzed (n=2). Yeast strains with genome integration (GI) and plasmid-based expression (PE) were included for all conditions. The YBR027C strain was added as an additional control since it was not predicted to be involved in methionine metabolism or methylation. Significance was determined using *t* tests with * indicating P ≤ 0.05, ** indicating P ≤ 0.01 and *** indicating P ≤ 0.001.

## Discussion

The present study is the first to prove that digitized mass fingerprints can be deciphered to rapidly predict genotypes and gene functions of unknown yeast knockouts. Despite the limitations of this proof-of-concept study, random forests and SVM algorithms were both effective at prediction GO accessions from the MALDI-TOF spectra of yeast gene knockout strains. Although random forests false negative rates were always perfect, there is a tradeoff with lower true positive prediction ability relative to SVM. Comparison of AUC scores emphasize that SVM is best the best overall GO prediction model, however random forests was better for 10 groups with large sample size (Figure 4). These results suggest that utilization of both SVM and random forests algorithms in a combined ensemble method might further improve GO accession prediction.

Machine learning prediction resulted in good coverage of smaller GO accession groups relative to *S. cerevisiae* genes and gene products listed in the AmiGO 2 database (http://amigo.geneontology.org/amigo). In this study, 75 binary vectors covering 45 genes were correctly predicted for GO:0005525 (GTP binding), which contains 161 *S. cerevisiae* genes in AmiGO 2 (28% coverage). Less specific GO accessions contained many genes that encode proteins larger than 20 kDa and coverage was lower for these cases. In the case of GO:0030529 (replaced by GO:1990904, ribonucleoprotein complex), a larger GO group with 868 listed *S. cerevisiae* genes in AmiGO 2, we identified 159 binary vectors covering 101 genes (11.6% coverage). For the very large GO accession GO:0008152 (metabolic process), binary vectors representing 244 genes were matched (5.7% coverage of 4,271 genes).

Data for each model was generated using a mass window of *m/z* 3,000 to 20,000, which should cover more proteins than metabolites; therefore, the current method may be primarily detecting differences at the proteomic level, rather than the metabolomic level. Accordingly, we hypothesize that this allows for prediction of gene functions for individual genes encoding proteins which produce ions of *m/z* 3,000 to 20,000. However, for strains with variations in multiple genes, it should be a greater challenge to predict multiple specific functions.

The current machine learning workflow was able to suggest the function of 28 uncharacterized genes, with metabolomics results consistent with the predictions for YDR215C and YLR122C. This workflow can be easily applied to additional strain libraries of various cell types, especially microbial production hosts. In the future, advanced artificial intelligence models should be developed to improve the prediction of specific gene functions from rapid mass fingerprints. The development of machine learning-based prediction methods is essential to realize the design, build, test and learn workflow of synthetic biology.

## Methods

### Cell extract preparation for MALDI-TOF analysis

The Yeast Deletion Mat-A Complete Set was obtained from Invitrogen. A replicator was used to inoculate glycerol stocks of 4,847 BY4741 *S. cerevisiae* gene knockouts into yeast extract peptone dextrose (YPD) medium in the wells of 96-well microplates with breathable sealing. After cultivation in YPD medium for 24 hours at 30 °C with shaking at 800 rpm, cells were pelleted by centrifugation and each well was washed two times with 180 μL Milli-Q water. The washed cell pellets were suspended in 70% formic acid (30 μL). The formic acid extracts were vacuum dried and resuspended in 15 μL Milli-Q water. After stirring and centrifugation, 1 μL of each supernatant was spotted onto MALDI plates and dried. 1 μL of matrix solution was then added and dried before MALDI-TOF mass analysis.

### MALDI-TOF yeast fingerprinting

Automatic high-throughput MALDI-TOF analysis was performed on a Bruker ultrafleXtreme operated in linear mode at 2 kHz. 384 spot MALDI plates were used for all experiments.

For clustering analysis, a total of 2,000 quality shots were obtained from each MALDI spot. For machine learning data collection, 25 MALDI-TOF shots were taken until a total of 200 quality shots with good signal to noise (S/N) data were obtained. If 200 quality shots (8 cycles) could not be obtained after 20 cycles of 25 shots, the spot was not used.

To collect the spectra for machine learning, all 4,847 gene knockout extracts were spotted in duplicate, resulting in 1,254 usable single spectra (for 1,254 knockouts) and 3,820 usable duplicate spectra (for 1,910 knockouts). To verify reproducibility, 74 gene knockouts were independently analyzed by a different operator resulting in 3 replicates for 25 genes, 4 replicates for 43 genes, 5 replicates for 1 gene and 6 replicates for 5 genes. All together, 3,238 gene knockouts and 5,356 high quality MALDI-TOF spectra were included in the analysis.

### Clustering analysis

MALDI-TOF spectra were digitized into 1,700 digit binary vectors by dividing a window of *m/z* 3,000 to 20,000 into 10 *m/z* intervals, followed by assigning a value of 0 for intervals with a standard score of maximum peak intensity under 52, or a value of 1 for intervals with a standard score of maximum peak intensity of 52 or higher. The distances between each binary vector were calculated by Euclidean distance, and binary vectors were clustered using Ward method. Correlation of each cluster with GO accessions was performed using the PANTHER Classification System (pantherdb.org).

### Scoring and machine learning

For computational prediction models, processed spectra were converted to binary vectors using the same method as that used for clustering analysis. Machine learning models were built using GO accessions associated with three or more binary vectors. This consisted of 4,731 binary vectors associated with 1,543 GO accessions for SVM, which was updated to 5,356 binary vectors representing 1,559 GO accessions for Tanimoto and random forests methods. Tanimoto, random forests and SVM prediction models were also evaluated based on 332 GO accessions matching 20 or more binary vectors.

For computational prediction, GO accession information was obtained from the AmiGo 2 database (http://amigo.geneontology.org/amigo). Positive and negative learning examples are defined as the 1,700-digit binary vectors either matching or not-matching a particular GO accession, respectively. In most cases, the number of positive examples was increased to 1,000 using SMOTE (synthetic minority over-sampling technique) and in a few cases where positive examples were more than or equal to 1,000, negative examples were increased to an amount 5-fold higher than positive examples.

Tanimoto scoring is a common method for comparing chemical similarity as utilized by the enzyme prediction software M-path (25, 26). The Tamimoto coefficient, an extension of the Jaccard coefficient, was employed to calculate similarity between binary vectors, using the following equation:

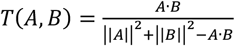

where *A* is a binary vector, *B* is a positive example average vector, and ||*X*|| represents the absolute value of *X*. Within each of the 1,559 GO accessions, Tanimoto coefficients were calculated for each binary vector by scoring against a positive example average vector.

The protocol for random forests prediction was similar to the described methods of Brieman (27). Eight variables were randomly selected from the dimensions of binary vectors, extracted from the same position of each binary vector, and used to create decision trees as weak learners for each GO accession group. This process is performed 500 times by randomly varying the 8 variables, resulting in 500 decision trees for each GO accession. The majority decision from the 500 trees is taken as the final random forests result.

SVM is another popular learning model for classification in which a boundary between positive negative examples is defined (28). SVM models were built based on the methods of Cortes and Vapnik, Bishop, and Chang and Lin (28, 32, 33). Positive examples and negative examples are mapped in high-dimensional feature space using a radial basis function (RBF) kernel. The models were built using soft margin SVM that allow for some misclassification. The SVM hyperparameters γ and C were optimized. As γ and C increase, learning data can be well discriminated, but overfitting will occur if γ and C becomes too large.

For the initial SVM models, cross validation tests were performed by building models with two-thirds of the training data, followed by testing the model with the remaining one-third of training data. This process was performed three times, with the one-third test data varying each time. For the Tanimoto and random forests predictions, cross validation was performed in the same manner except using a 10-fold process rather than the above described 3-fold process. 1,543 GO accessions matching 3 or more binary vectors were used for initial SVM models, and 1,559 GO accessions matching 3 or more binary vectors were used for Tanimoto scoring and random forests prediction. 332 GO accessions matching 20 or more binary vectors were used for Tanimoto, random forests and SVM prediction.

True positive rates were calculated by dividing the number of true positives by the sum of the true positives and false negatives, while true negative rates were calculated by diving the number of true negatives by the sum of true negatives and false positives. False negative rates were calculated by dividing the number of false negatives by the sum of the false negatives and true positives, while false positive rates were calculated by diving the number of false positives by the sum of false positives and true negatives.

### Testing spectra from knockout strains of genes with unknown function

69 gene knockouts corresponding to 110 vectors were identified as genes with unknown function according to the AmiGO2 database. The 110 vectors of unknown gene knockouts were therefore used as test samples, in order to obtain predicted functions for each gene.

For prediction of unknown gene functions, SVM models were built using the set of 5,356 MALDI-TOF spectra-derived binary vectors. Models were created for the 332 GO accessions, which matched to 20 or more binary vectors. Accuracies of the models were then verified by 10 cross-validations. For GO prediction of a knockout strain of an unknown gene, scores are given as an average of results from 10 models obtained by 10 cross-validations.

In the current study, functions were predicted for 41 knockouts of uncharacterized genes with duplicate MALDI-TOF spectra. If both mass fingerprint-derived binary vectors for an unknown knockout showed positive matching to a GO accession with less than 400 positive training vectors, then this was considered as a positive match. Average SVM scores obtained from two mass spectra-derived binary vectors are presented in Figure 7.

### Metabolomics analysis of yeast strains

Yeast strains were precultured in YPD and then transferred to minimal medium (containing 6.7 g / L yeast nitrogen base, 2% glucose, 21 mg / L histidine, 120 mg / L leucine, 60 mg / L lysine, 20 mg / L tryptophan, 20 mg / L arginine, 20 mg / L tyrosine, 40 mg / L threonine, 50 mg / L phenylalanine, 20 mg / L uracil and 20 mg / L adenine), with matching initial cell densities. Cultures were then grown at 30 **°**C while shaking at 150 rpm. Approximately 24 hours later, 5 mL of each yeast culture was added to 7 mL methanol chilled at -30 **°**C; the quenched samples were centrifuged and processed for metabolite extraction according to our previous reports (34). Intracellular metabolites were then quantified by liquid chromatography-mass spectrometry (LC-MS) on a Shimadzu LCMS-8050 system as described in our previous reports (35). Metabolomics results were analyzed with Shimadzu LabSolutions and Prism 7.

### Conversion of norcoclaurine to coclaurine and *N*-methylcoclaurine in yeast

*Papaver somniferum* norcoclaurine 6-*O*-methyltransferase (6OMT) and coclaurine *N*-methyltransferasea (CNMT) genes were introduced into yeast strains according to the methods of Ishii *et al*. (36). Vectors pATP405red-optPs6OMT-optPsCNMT (genome integration vector) and pATP425-optPs6OMT-optPsCNMT (plasmid type vector) were constructed with *S. cerevisiae* codon optimized methyltransferase genes.

Yeast strains with matching initial cell densities were grown in 2.5 mL minimal medium (containing 6.7 g / L yeast nitrogen base, 2% glucose, 20 mg / L histidine, 20 mg / L uracil and 15 mg / L methionine) at 30**°**C with shaking at 200 rpm. Leucine (60 mg / L) was included in the minimal medium for the wild-type strain, but two knockout strains were also tested with the leucine-containing medium and no effect was found on the growth. 105 µL of 25 mM norcoclaurine (1 mM final concentration) was added to each yeast culture. A wild-type pATP405red-optPs6OMT-optPsCNMT control culture with water added in place of norcoclaurine was included. After conversion of norcoclaurine for approximately 60 hours, 400 µL of each culture was collected and filtered using Millipore Amicon Ultra-0.5 mL centrifugal filters, and the filtered samples were stored at -80 **°**C. Benzylisoquinoline alkaloid content of supernatants were then quantified by LC-MS on a Shimadzu LCMS-8050 system as described in our previous reports (29, 30). Results were analyzed with Shimadzu LabSolutions and Prism 7.

## Author Contributions

MA, KH, AK, TH and CJV conceived and designed the experiments. CJV, MM, TY, MM, MN, JI and KH performed the experiments. CJV, MM, SY, TY, MM, NW, JI, HSA, and MA analyzed the data. CJV, SY, HSA and MA prepared the manuscript.

## Acknowledgements

The authors thank Ryo Suzuki and Tomomi Nakamura for their help with construction and cultivation of benzylisoquinoline alkaloid producing yeast strains. The research in this manuscript was funded by project P16009, Development of Production Techniques for Highly Functional Biomaterials Using Smart Cells of Plants and Other Organisms (Smart Cell Project) from the New Energy and Industrial Technology Development Organization (NEDO), and The Program for Forming Japan’s Peak Research Universities (J-PEAKS) from the Japan Society for the Promotion of Science (JSPS). Machine learning research of CJV is supported by the G-7 Scholarship Foundation and the Takeda Science Foundation.

## Supplementary Information

### Clustering mass-based vectors for GO prediction

For our first attempt using clustering analysis, the 4,847 knockouts of the yeast library were refined to a much smaller set that could be better representative of the *m/z* 3,000-20,000 spectra. Genes encoding proteins with a molecular weight of 20 kDa or more (4,961 genes) were removed resulting in 939 genes. After removal of hypothetical, ribosomal and un-annotated genes, this was further narrowed down to 103 genes of which 74 corresponding gene knockouts were present in the yeast knockout library. The selected 74 genes were clustered into 13 groups as displayed in Supplementary Figure 1.

Each cluster was correlated with matching GO accessions using the PANTHER Classification System (pantherdb.org). Clusters 3-4 contained mostly wildtype binary vectors. Although GO correlation could be enriched from the clustering analysis for knockouts of genes encoding proteins of 20 kDa or less, this strategy was too simple for analysis of the entire set of 3,238 gene knockouts (5,356 spectra). To extend the GO correlation to the entire set of binary vectors, Tanimoto scoring, and machine learning algorithms were explored (Figure 3, lower path).

**Supplementary Figure 1.**
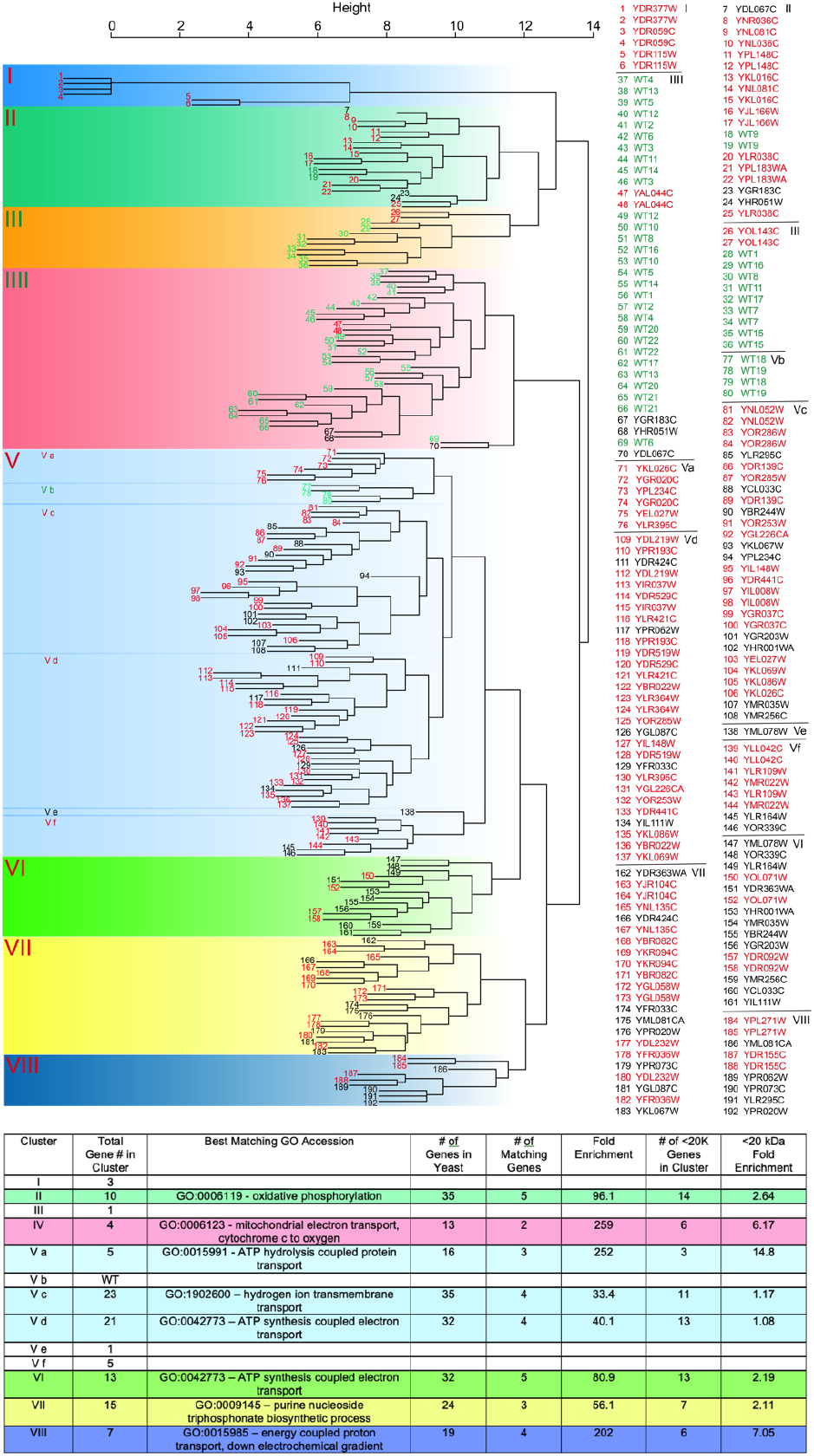
GO analysis of yeast knockout derived binary vector clusters. In the upper tree, wild-type and knockout duplicates are colored green and red, respectively, when they appear in the same cluster. Yeast knockout binary vectors with duplicates in a separate cluster are colored black. The clustering analysis was performed with 74 knockout spectra in duplicate and 44 total wild-type spectra. In the lower table, the best GO accession matches are listed for each cluster. For clusters I, III, V b, V e and V f, there was no result due to small gene sample size. Fold enrichment is calculated with the following equation: (number of matching genes / number of genes in cluster) / (total number of yeast genes matching the GO accession / total number of yeast genes [6,728]). Fold enrichment for genes encoding proteins under 20 kDa was calculated using the same equation using the number of genes encoding proteins under 20 kDa in place of total gene number.

**Supplementary Figure 2.**
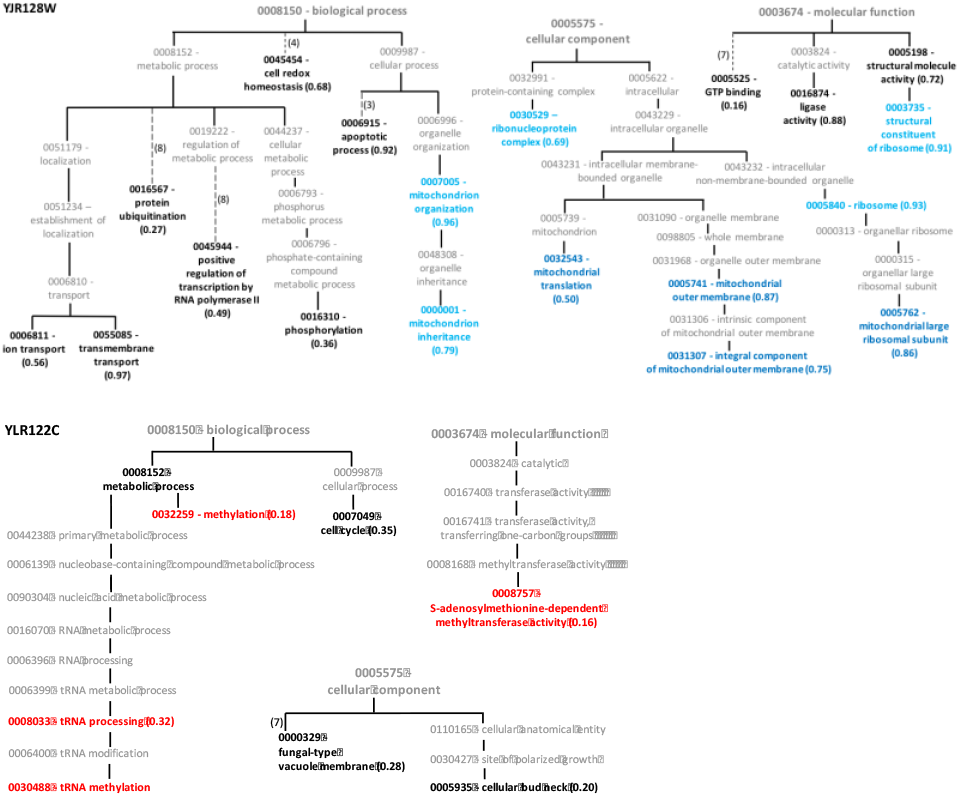
Examples of GO matching to yeast knockouts of genes with unknown functions. Matching GO accessions are shown in bold.

